# EZH2 inhibition results in genome-wide PRC2 redistribution

**DOI:** 10.1101/713842

**Authors:** Chih-Chi Yuan, Ah Jung Jeon, Greg Tucker-Kellogg, Barbara Bryant, Patrick Trojer

**Affiliations:** Constellation Pharmaceuticals, Cambridge, Massachusetts, 02142, USA; Department of Biological Sciences, National University of Singapore, Singapore 117558, Singapore; Computational Biology Programme, Faculty of Science, National University of Singapore, Singapore 117558, Singapore

**Keywords:** EZH2, PRC2, Lymphoma, DLBCL, EZH2 inhibition

## Abstract

Histone methyltransferase polycomb repressive complex 2 (PRC2) plays a critical role in cell fate determination, and its catalytic subunit EZH2 is a key oncogenic driver in GCB-DLBCL. EZH2 inhibition in some GCB-DLBCL cell models leads to a global loss of H3K27me3, the derepression of a subset of silenced PRC2 target genes, and ultimately cell death. Here we show that EZH2 inhibition causes global redistribution of PRC2 components. We observe a reduction in the already-low levels of PRC2 at active genes. On the other hand, focal PRC2 accumulation and concomitant H3K27me3 retention occur at many canonical embryonic stem cell PRC2 nucleation sites. PRC2 accumulation is also enriched in plasma/memory cell genes repressed by PRC2 activity in the germinal center. We see PRC2 redistribution to, and H3K27me3 retention at, differentiation-related genes not only in cell lines that are insensitive to killing by EZH2 inhibition, but also in sensitive cell lines. Thus, PRC2 redistribution to B cell differentiation genes is not sufficient to explain the resistance to EZH2 inhibitors in some DLBCL cell lines.

## Introduction

Research over the past several years has demonstrated that chromatin regulating factors are frequently targeted in tumorigenesis (Flavahan et al 2017; Morgan and Shilatifard 2015). Polycomb repressive complex 2 (PRC2) is one of the most investigated chromatin regulating factors. Both gain and loss of function mutations in PRC2 components have been implicated in cancer formation (Helin 2016). PRC2 is one of several Polycomb group (PcG) protein complexes and plays an important role in body plan formation during animal development by regulating the expression of homeobox genes at different body segments (Margueron and Reinberg 2011). PcG proteins are also critical in maintaining cellular memory of cell identity in later stages of development by repressing differentiation toward other cell types (Lee et al. 2006; Thornton et al. 2014). PRC2 is a transcription repressive factor composed of core components EZH2, SUZ12, and EED. At the molecular level, PRC2 catalyzes the mono-, di-, and tri-methylation of histone H3K27 (Margueron and Reinberg, 2011) and H3K27me3 is the hallmark of facultative heterochromatin and transcription silencing (Di Croce and Helin, 2013; Kirmizis et al., 2004; Margueron and Reinberg, 2011). PRC2 represses transcription by methylating H3K27 at its target genes and induces the formation of repressive chromatin structure.

In a number of cancers, the EZH2 gene is often either amplified, overexpressed, or carries an activating mutation that alters its substrate specificity (Hock, 2012); cancer cells use the altered chromatin landscape and transcription output from the abnormal PRC2 activities to gain survival advantage. In particular, tumors from a high percentage of germinal center B cell-like diffuse large B cell lymphoma (GCB-DLBCL) patients carry a somatic mutation within EZH2 that promotes the conversion of H3K27me2 to H3K27me3 (Morin et al., 2010). DLBCL is the most common type of non-Hodgkin’s lymphoma, but it is heterogeneous; GCB-DLBCL is a molecularly defined subtype of DLBCL in which the tumor cells are thought to arise from germinal center B cells (Lenz et al., 2008). EZH2 activity in the germinal center during normal B cell development temporarily suppresses genes whose activity is restored upon terminal differentiation and plasma/memory cell formation. It has been proposed that the EZH2 activating mutation in GCB-DLBCL leads to the sustained hypermethylation of those loci, forcing the terminal differentiation genes deeper into a repressive state, thus preventing both their reactivation and terminal B cell differentiation (Béguelin et al., 2013).

In the face of accumulating preclinical evidence that EZH2 plays a critical role in the development of several cancer types including GCB-DLBCL and prostate cancer, EZH2 inhibitors have been developed and several are currently in clinical trials in different disease indications (Gulati et al., 2018; Italiano et al., 2018; Yap et al., 2016). EZH2 inhibition leads to a global reduction of H3K27me3 and eventually cell death in a dose and time-dependent manner in many cancer cell models (Bradley et al., 2014; Garapaty-Rao et al., 2013; Girard et al., 2014; McCabe et al., 2012; Tiffen et al., 2015; Zeng et al., 2017)

While EZH2 inhibition leads to global reduction of H3K27me3, recent research has indicated that EZH2 inhibition does not reduce H3K27me3 levels equally across the genome (Xu et al., 2015). In this study, we confirm and extend these observations, showing that PRC2 components are redistributed to a number of loci that also retain relatively higher levels of H3K27me3. We observe redistribution to at least two categories of genes: previously observed H3K27me3 and PRC2 loci in embryonic stem cells, and germinal center B cell terminal differentiation genes normally repressed by PRC2 in GCB-DLBCL. Because re-expression of the GCB differentiation genes has been proposed as a mechanism of action for EZH2 inhibitors in GCB-DLBCL, their continued repression as a result of PRC2 accumulation is a candidate mechanism for resistance to EZH2 inhibitors. However, as we see redistribution of PRC2 and retention of repressive H3K27me3 at terminal differentiation genes in both EZH2 inhibitor insensitive and sensitive cell models, our data suggest that accumulation of PRC2 on differentiation genes is not sufficient to explain resistance to EZH2 inhibitors.

## Results

### PRC2 accumulates at H3K27me3-retaining TSSs and is lost from active TSSs upon EZH2 inhibition

We previously described the characterization of two small molecule inhibitors, CPI-360 and CPI-169, of EZH2 which showed time- and dose-dependent efficacy in Non-Hodgkin’s Lymphoma models in vitro and in vivo (Bradley et al., 2014). These EZH2 inhibitors demonstrated robust target engagement in cell models by decreasing global H3K27me3 and H3K27me2, but not H3K27me1 levels (Bradley et al., 2014). To assess the effect of EZH2 inhibition on H3K27me3 and PRC2 distribution across the genome, we treated KARPAS-422, a DLBCL cell model that is sensitive to EZH2 inhibitor treatment, with DMSO control or EZH2 inhibitor for 4 and 8 days and carried out ChIP-seq campaigns for histone marks H3K27me3 and H3K4me3, and PRC2 components EZH2 and SUZ12.

H3K27me3 is globally reduced upon multi-day EZH2 inhibition while PRC2 components EZH2 and SUZ12 remain unchanged. This is shown for CPI-169 in Figure S1A, and in Figure 2B of Bradley 2014 for CPI-360. Analysis of ChIP-seq data reveals that in DMSO-treated cells, H3K27me3 is abundant and occupies broad segments of the KARPAS-422 genome. EZH2 inhibition greatly reduces H3K27me3 levels but not PRC2 components EZH2 and SUZ12 on chromatin (see examples in Figure S1B). On the other hand, H3K4me3 levels remain steady both at the bulk level (Figure S1A) and at randomly selected genomic regions (Figure S2). Because promoter-associated H3K27me3 is associated with repression of gene expression, we examined the ChIP-seq signal around transcription start sites (TSSs). The average ChIP-seq signal at TSSs was consistent with the Western blot observations: H3K27me3 signal showed marked reduction but EZH2, SUZ12 and H3K4me3 showed little change (Figure S1C).

Although the level of H3K27me3 was decreased at most loci, several distinct patterns of H3K27me3 distribution were observed around TSSs. To explore these patterns, we performed k-means clustering of the H3K27me3 signal at TSS regions in both DMSO- and EZH2 inhibitor-treated cells (Figure 1A). We found that although the level of H3K27me3 was reduced in all clusters, not all H3K27me3-marked TSSs responded to EZH2 inhibition the same way: cluster 1, for example, retained much higher levels of H3K27me3 after inhibitor treatment than all other clusters. Interestingly, cluster 1 also showed the highest average H3K27me3 signal at baseline, suggesting that retention of H3K27me3 preferentially occurs at high H3K27me3 sites.

**Figure 1.**
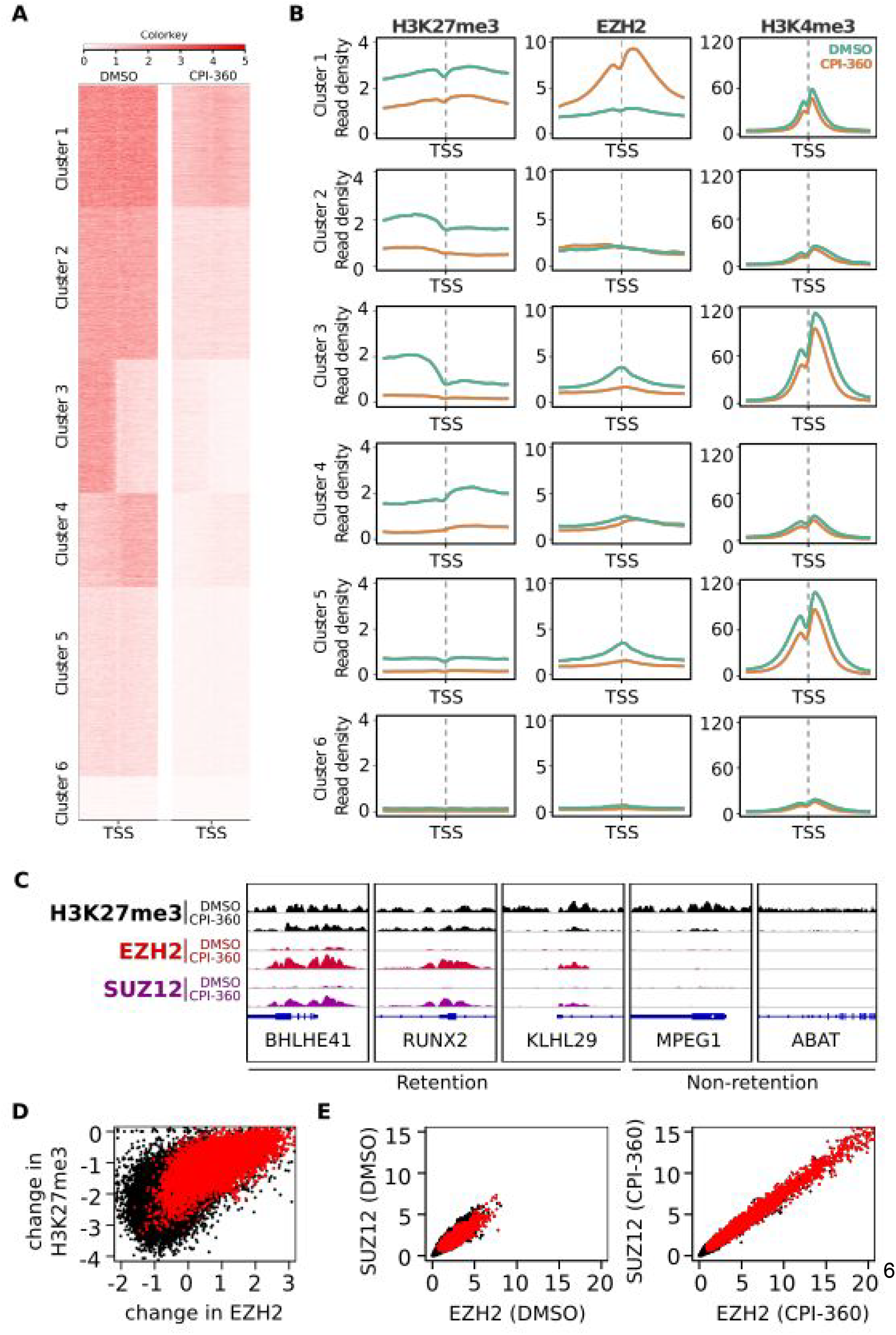
Differential patterns of H3K27me3 at TSSs and H3K27me3 retention phenomenon. A. Heatmaps of the six H3K27me3 clusters at TSS regions (+/− 1.5kb) reveal different patterns of mark changes upon EZH2 inhibition in KARPAS-422 cells. B. Average H3K27me3, EZH2, and H3K4me3 signal for each TSS cluster, with and without EZH2 inhibition, for the region +/− 1.5kb of the TSS. TSSs in cluster 1 show increased EZH2 signal upon EZH2 inhibitor treatment. C. ChIP-seq signal for H3K27me3, EZH2, and SUZ12 are shown for representative retention and non-retention TSSs. D. Change in H3K27me3 vs change in EZH2 signal across all TSSs. Red indicates TSSs from cluster 1. Change is regularized log2 fold change. TSSs that accumulate EZH2 tend to retain more H3K27me3. E. Scatter plots of ChIP-seq signal downstream of each TSS for EZH2 and SUZ12, treated with DMSO or CPI-360. Red indicates TSSs in cluster 1. EZH2 and SUZ12 are strongly correlated both at baseline and after EZH2 inhibitor treatment.

To investigate the H3K27me3 retention phenomenon further, we determined the average H3K27me3, EZH2 and H3K4me3 levels within each H3K27me3 TSS cluster (Figure 1B). H3K4me3 is highest in clusters 3 and 5, where H3K27me3 levels are relatively low downstream of the TSS, suggesting that these clusters comprise actively transcribing genes. H3K4me3 levels showed little change upon EZH2 inhibition, consistent with the average signals from aggregated TSSs (Figure S1C), and the Western blot (Figure S1A). Unexpectedly, EZH2 patterns changed significantly, with redistribution away from TSSs in clusters 3 and 5, and strong gain of signal in cluster 1 (Figure 1B, middle column). The results suggest that the retention of H3K27me3 is associated with a marked accumulation of chromatin-bound EZH2. Examples of H3K27me3 retention and non-retention loci are shown in Figure 1C.

To address whether the correlation between EZH2 accumulation and H3K27me3 retention held true at individual transcript start sites, we plotted the H3K27me3 change upon EZH2 inhibition against the change in EZH2 signal for each TSS. The resulting scatter plots show that cluster 1 TSSs (red) contained most of the TSSs with an accumulation of EZH2 and the least reduction of H3K27me3 (Figure 1D), while TSSs from other clusters showed greater H3K27me3 loss and no change or a loss of EZH2 (Figure S3). Furthermore, a scatter plot of baseline H3K27me3 against relative change in EZH2 signal demonstrates that TSSs with high baseline H3K27me3 are more likely to accumulate EZH2, and most of these TSSs are within cluster 1 (Figure S4). These results reiterate the tendency that upon EZH2 inhibition, EZH2 accumulation preferentially occurs at TSSs with higher H3K27me3 levels. H3K27me3 retention and EZH2 accumulation were confirmed with ChIP-qPCR at three retention loci (BHLHE41, KLHL29, and RUNX2) and two non-retention loci (ABAT1, MPEG1, Figure S5).

To determine whether EZH2 accumulates at TSSs on its own or as part of the PRC2 complex, SUZ12 signal intensities were plotted against EZH2 for each TSS (Figure 1E). The occupancy levels of the two proteins at TSSs are highly correlated at both baseline (R=0.842) and upon EZH2 inhibition (R=0.961), and both proteins accumulate on cluster 1 TSSs upon EZH2 inhibition (Figure 1E, right panel). The strong correlation between EZH2 and SUZ12 binding is consistent with a tight functional link between the two proteins and suggests that the PRC2 complex is redistributed throughout the genome in response to EZH2 inhibition.

### PRC2 preferentially redistributes to TSSs and CpG islands upon EZH2 inhibition

The striking accumulation of EZH2 and SUZ12 at a subset of TSSs upon EZH2 inhibition prompted us to investigate whether similar accumulation occurs at other genomic elements upon EZH2 inhibitor treatment. CpG islands and active enhancers were selected for the analysis due to the critical roles of these elements in transcription regulation. Active enhancers were identified as H3K27ac-positive regions. Interestingly, we saw a strong average increase of EZH2 signal at CpG islands and a strong loss of EZH2 at H3K27ac sites in EZH2 inhibitor treated cells (Figure 2A, top panels). These results demonstrate that EZH2 accumulation occurs preferentially at a subset of genomic elements upon EZH2 inhibitor treatment. SUZ12 showed similar changes at these genomic elements (Figure 2A, lower panels), suggesting a global redistribution of PRC2 complex upon EZH2 inhibitor treatment.

**Figure 2.**
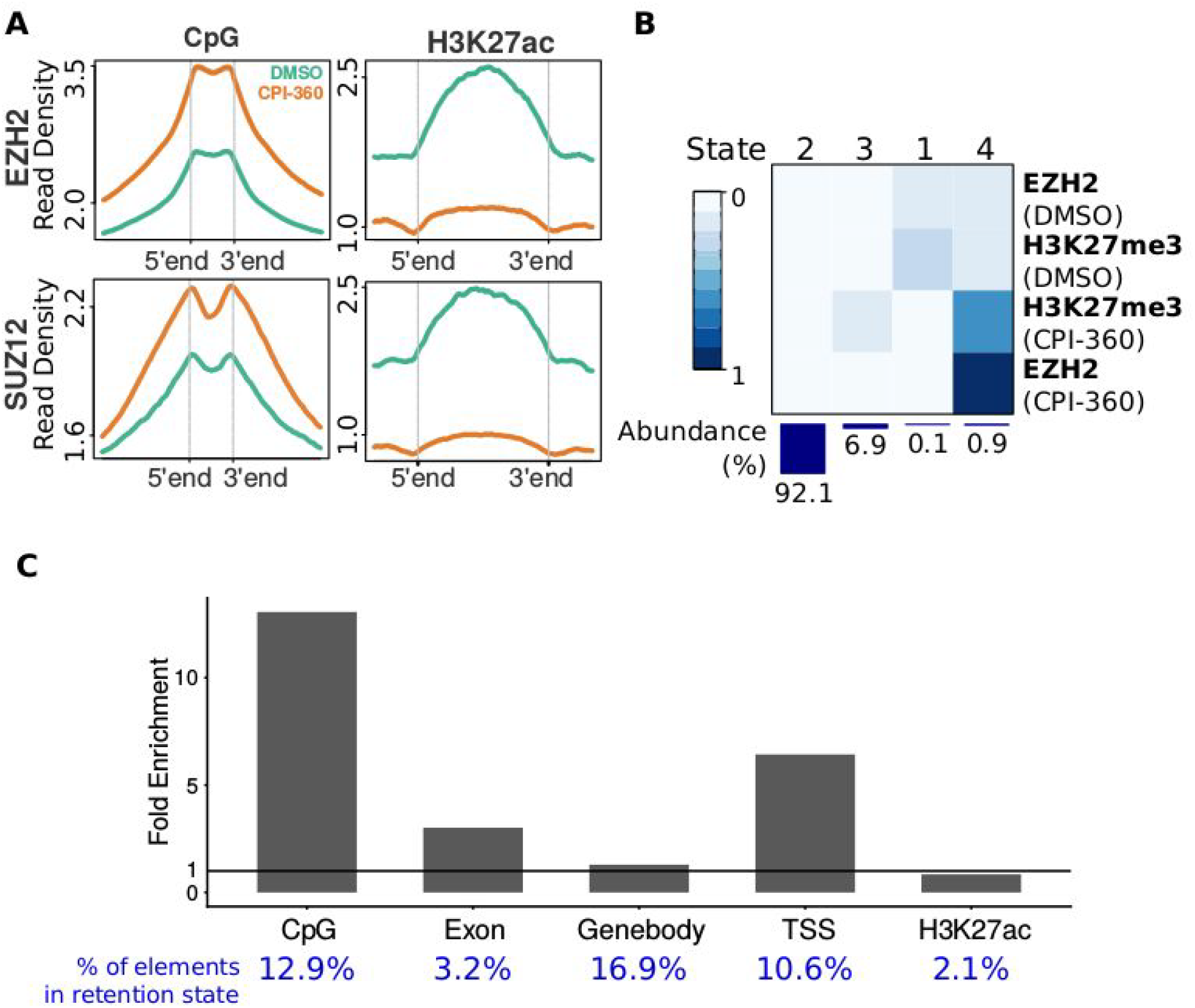
EZH2 redistribution upon EZH2 inhibitor treatment and retention site analysis. A. Gain of EZH2 and SUZ12 at CpG islands and loss of EZH2 and SUZ12 at H3K27ac loci upon EZH2 inhibition. B. Emission probability matrix for ChromHMM-based definition of chromatin states. For further analysis, EZH2 retention was defined as genomic regions in State 4. The bar plot shows the relative abundance of each chromatin state in the genome. C. Enrichment of retention state at genomic elements relative to average across the genome. The fraction of each type of genomic element having retention state is shown in blue.

To investigate PRC2 accumulation and H3K27me3 retention genome-wide, the ChIP-seq datasets were analyzed with the chromatin state calling algorithm ChromHMM (Ernst and Kellis, 2012). We defined retention sites as those genomic loci with a chromatin state having high probability of EZH2 and H3K27me3 in the EZH2 inhibitor treated sample (Figure 2B). This definition identified 21,112 contiguous segments (144,720 200 bp segments) covering roughly 0.92% of the genome as being in the retention state in EZH2 inhibitor treated KARPAS-422 cells.

We tested for overlap of the retention state segments with various types of genomic elements, and confirmed that retention sites were highly enriched in CpG islands and TSSs relative to the rest of the genome (Figure 2C). Gene bodies and regions marked with H3K27ac were not enriched, while exons showed slight enrichment. The presence of a TSS or CpG island, however, was not sufficient to establish a retention site: only 12.9% of CpG islands and 10.6% of TSSs overlapped at all with a retention site.

### H3K27me3 retention sites are partially conserved across cancer cell models

The observed H3K27me3 retention and PRC2 redistribution in KARPAS-422 upon EZH2 inhibition raises the question of whether the retention of H3K27me3 and the redistribution of PRC2 is locus-specific, and whether it is common among cancer cell lines. To answer this question, we studied the distribution of H3K27me3 and EZH2 in 5 additional cell models: two DLBCL cell lines with activating EZH2 mutations (RL and SU-DHL-6) and three multiple myeloma cell lines (KMS28BM, KMS28PE, and RPMI8226) with wild type EZH2. ChromHMM analysis identified H3K27me3 retention sites upon EZH2 inhibition in all 5 cell models, suggesting H3K27me3 retention is a common phenomenon. Consistent with the observations in KARPAS-422 (which also has an activating EZH2 mutation), the retention sites in all cell lines are enriched in TSSs and CpG islands (Figure 3A). Furthermore, many retention sites are conserved across cell lines, while others are found only in a single cell line (see examples in Figure S7A).

**Figure 3.**
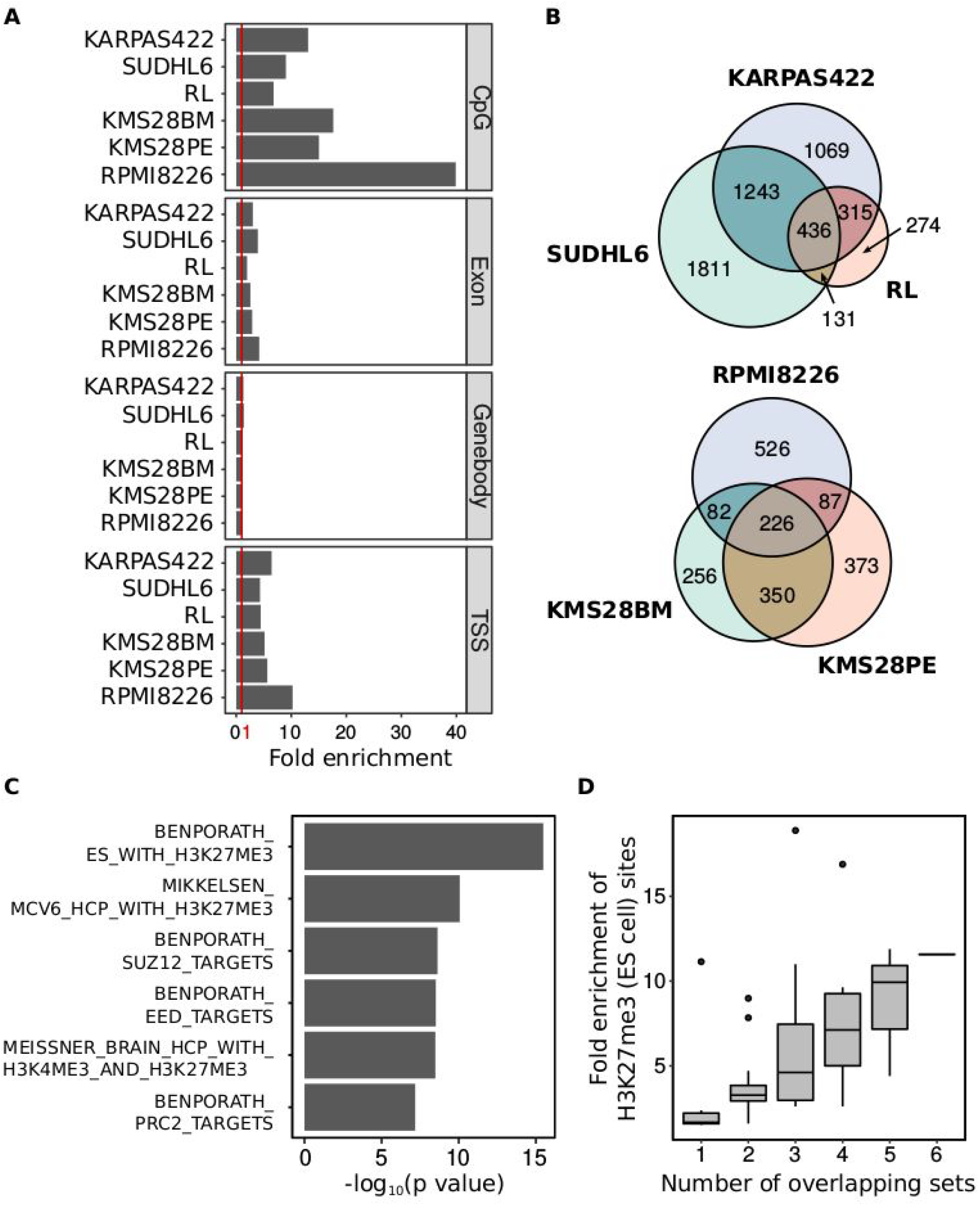
Retention site in cells from different origins are partially conserved. A. Fold enrichment of retention sites for different genomic features across different cell lines. CpG islands and TSS regions are consistently enriched, though the total enrichment and number of elements with retention varies between cell lines. B. Overlap of retention genes across three DLBCL (KARPAS422, SUDHL6, RL) and three multiple myeloma cell models (RPMI8226, KMS28BM, KMS28PE). C. Functional analysis (MSigDB) of 55 retention genes conserved across all six cell lines D. Functional analysis showing more conserved retention genes have stronger enrichment of the BENPORATH_ES_WITH_H3K27ME3 gene set.

To further examine the conservation of H3K27me3 retention and PRC2 accumulation across cell lines, we defined retention TSSs as TSSs within 2.5 kb of an H3K27me3 retention site. In KARPAS-422, 68% of retention TSSs are in cluster 1 and 22% are in cluster 2. Of the 5,379 retention TSSs observed in the three lymphoma cell lines, 436 are conserved, and 226 out of the 1,900 retention TSSs in the myeloma lines are common between the three cell lines (Figure 3B, S8). Overall, 55 TSSs are conserved among all six cell lines (Figure S7B). The conservation of retention sites across cell lines occurs more frequently than would be expected at random. Moreover, the better-conserved retention sites showed higher enrichment over random background (Figure S8C). Interestingly, however, fewer than 50% of the retention TSSs are conserved between KMS28BM and KMS28PE, two multiple myeloma cell lines derived from the same patient, suggesting that the formation of retention sites can also be dynamic as cancer cells evolve. MSIGDB analysis (Liberzon et al., 2011) of the 55 common retention TSSs showed a very strong overlap with PRC2 binding sites in embryonic stem cells (Figure 3C), and the more conserved a retention TSS, the more likely that TSS is to be a PRC2 target sites in embryonic stem (ES) cells (Figure 3D, S7B).

### Many H3K27me3 retention TSSs are also marked with H3K4me3

Transcription start site regions can be monovalent — marked with either repressive H3K27me3 or activation-associated H3K4me3 — or in a bivalent state carrying both marks (Bernstein et al., 2006) Comparing H3K27me3 and H3K4me3 ChIP-seq signals in KARPAS-422, we found that some of the retention-site-containing TSSs are bivalent (Figure 4A). To examine co-occurrence of H3K27me3 and H3K4me3, H3K4me3 signal was plotted against H3K27me3 signal immediately downstream of the TSS (Figure 4B, retention genes in red): 34.8% of retention TSSs were bivalent, having average H3K4me3 and average H3K27me3 signals above our defined cutoffs.

**Figure 4.**
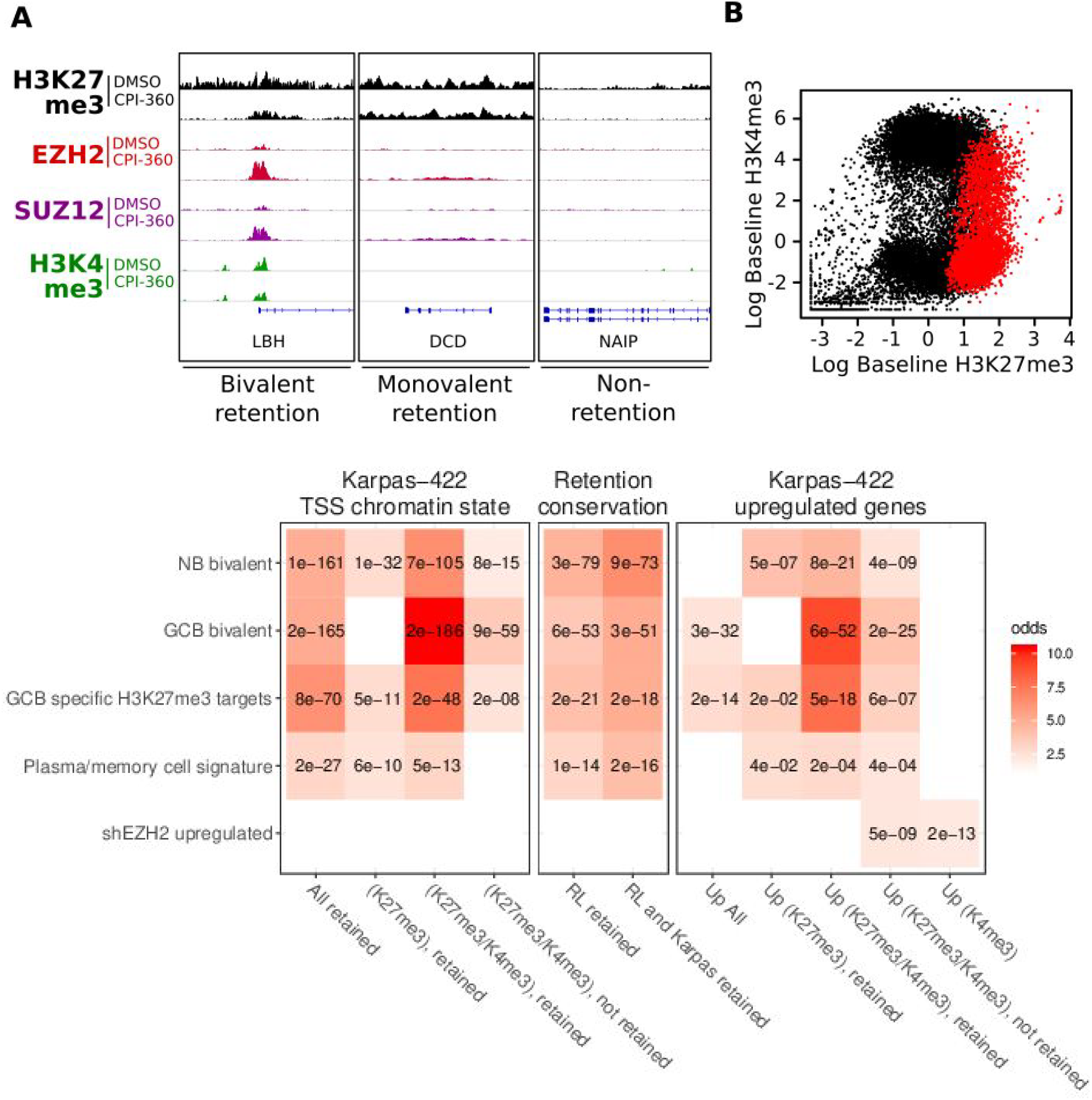
Retention genes from KARPAS-422 cells are enriched with GCB bivalent and plasma/memory cell signature genes. A. ChIP-seq tracks showing examples of non-retention, bivalent retention and monovalent retention genes. B. Many retention TSSs (red) are bivalent, having both H3K27me3 and H3K4me3 in DMSO-treated KARPAS-422 cells. C. Heatmaps of KARPAS-422 and RL retention genes showing enrichment of GCB bivalent and plasma/memory cell signature genes

### Gene expression changes are not limited to H3K27me3-repressed genes

Using previously published RNA-seq data from KARPAS-422 cells treated with DMSO or EZH2 inhibitor (GSE62056, Bradley et al., 2014), we examined gene expression levels for each cluster (Figure S6A). Clusters most highly marked with H3K27me3 had the lowest gene expression, while clusters with the greatest H3K4me3 levels showed the highest gene expression.

EZH2 inhibition leads to up- and down-regulation of many genes, as has been reported previously (Bradley et al., 2014). Up-regulation occurs in genes from all clusters (Figure S6B and C). Down-regulation, on the other hand, is much more prevalent in the actively transcribed genes of clusters 3 and 5 (Figure S6B and C), having high H3K4me3 and low H3K27me3. The top PRC2 recruitment genes are nearly all in cluster 1, but more of them are expressed, and more down-regulated by EZH2 inhibition, than other cluster 1 genes (data not shown). Bivalent (H3K27me3 + H3K4me3) genes are approximately three times more likely to be up-regulated than those marked by only H3K4me3 or H3K27me3 (Table S4). Overall, the patterns of gene expression change show that the transcriptional effects of EZH2 inhibition are not limited to the upregulation of genes directly silenced by H3K27me3.

### Germinal Center B-cell H3K4me3/H3K27me3 bivalent genes are highly enriched in retention genes

To explore the function of the retention genes, we performed gene set enrichment analysis against the MSIGDB C2 collection (Liberzon et al., 2011; Subramanian et al., 2005), and identified conserved H3K27me3 and PRC2 component target genes in ES cells as enriched and conserved across all cell lines (Figure 3C, D). Recent studies in GCB-DLBCL have identified additional gene sets that include GCB-specific H3K27me3 and bivalent targets, genes upregulated on EZH2 inhibition or knockdown, and genes upregulated on exit from the germinal center to form plasma/memory B cells (Béguelin et al., 2013) as well as H3K4me3/H3K27me3 bivalent genes in germinal center B cells (Béguelin et al., 2016). We tested for enrichment of these categories among retention genes in different chromatin states (Figure 4C, left), and identified high enrichment across multiple categories, with the highest enrichment for GCB-specific bivalent genes in our bivalent retained genes. Retained and bivalent retained genes also showed strong enrichment for plasma/memory cell signature genes and naive B cell bivalent genes, but not genes upregulated on EZH2 knockdown. The enrichment for GCB bivalent gene and plasma/memory cell signature genes is of particular interest, as it has been proposed that cells undergoing the germinal center reaction temporarily repress genes required for terminal differentiation by means of PRC2-mediated H3K27me3 addition to their promoters. Cancer cells harboring EZH2 activating mutations would hyper-repress these genes, thereby preventing the cells from exiting the proliferative state. The significant overlap between the KARPAS-422 bivalent retention genes and the GCB bivalent genes confirms the previously reported chromatin state at these genes in DLBCL, but also suggests that their hyper-repression in DLBCL is not readily lifted by EZH2 inhibition. Similarly, the enrichment of the plasma/memory cell signature amongst retention genes confirms previous reports of PRC2-mediated suppression of terminal differentiation in DLBCL, while suggesting a resistance to removal of the H3K27me3 mark in response to EZH2 inhibition.

### Enrichment of GCB-bivalent and plasma/memory cell signature genes is conserved between sensitive and insensitive DLBCL models

As there is significant overlap between retention genes and the plasma/memory cell signature in KARPAS-422, we asked whether this is a common phenomenon in other DLBCL cell lines. RL is an EZH2-mutant carrying DLBCL cell line that is insensitive to EZH2 inhibition; interestingly, PRC2 redistribution and H3K27me3 retention were still observed in RL upon EZH2 inhibition, and about 65% (751/1156) of the retention genes in RL are also retention genes in KARPAS-422 (Figure 3C). As in KARPAS-422, retention genes in RL show significant overlap with both the plasma/memory cell signature and with GCB bivalent genes (Figure 4C, middle). In fact, genes that are retention genes in both RL and KARPAS-422 also show significant overlap with both the differentiation and GCB bivalent signatures. That both of these EZH2-mutant cell lines repress these genes with H3K27me3 is consistent with germinal cell biology and the model of hyper-repression in DLBCL. However, EZH2 inhibition kills KARPAS-422 cells but not RL cells. The similarity of PRC2 accumulation and H3K27me3 retention between these cell lines suggests that de-repression of H3K27me3-marked differentiation genes does not fully explain sensitivity to cell killing.

## Discussion

### Retention of H3K27me3 and redistribution of PRC2

In this study we examined genome-wide changes in H3K27me3 and PRC2 upon EZH2 inhibitor treatment of cancer cell models. Consistent with previous data, we observed global loss of H3K27me3 upon inhibitor treatment; however, at a small subset of genomic loci a significant fraction of H3K27me3 is retained, accompanied by accumulation of PRC2. We also found that H3K27me3 retention and PRC2 accumulation tends to occur at genomic loci with high H3K27me3 level at baseline. Focal H3K27me3 retention has been reported in a variety of contexts, including EZH2 inhibition (Xu et al., 2015), deletion of EZH2 in a mouse model, and up-regulation of NSD2 (Popovic et al., 2014). Accumulation of PRC2 components has been previously reported upon NSD2 up-regulation (Popovic et al., 2014), mutation of H3.3K27 (Chan et al., 2013), and epithelial-mesenchymal transition (Malouf et al., 2013). Here, we report a striking level of focal PRC2 accumulation in the context of EZH2 inhibition.

It has been hypothesized that Polycomb-mediated repressive domains in the genome are formed by the spreading of PRC2 from nucleation sites at CpG islands where there is no DNA methylation and no transcription (Cooper et al., 2014; van Heeringen et al., 2014; Ku et al., 2008; Laugesen et al., 2019; Oksuz et al., 2018) Once recruited to the nucleation sites, PRC2 trimethylates H3K27 in the adjacent nucleosome, and the newly deposited H3K27me3 recruits PRC2 through interaction with the EED component of PRC2. The repeated PRC2 recruitment and H3K27me3 catalysis leads to spreading and establishing of polycomb-mediated repressive domains. Recently, Oksuz et. al. demonstrated the spreading of PRC2 from nucleation sites in a time dependent manner (Oksuz et al., 2018). The same mechanism of PRC2 recruitment to H3K27me3 and methylation of an adjacent histone tail could support maintenance of H3K27me3 domains in response to dilution with new nucleosomes during genome replication, or even removal of the histone mark by enzymatic demethylation. We postulate that during EZH2 inhibitor treatment, as the deposition of H3K27me3 is inhibited, these H3K27me3 domains can no longer be maintained across cell divisions. However, the H3K27me3-independent recruitment of PRC2 to the nucleation sites is still in effect. With the same amount of PRC2 available in the nucleus, and reduced binding sites available, PRC2 may accumulate on the remaining loci. This is similar to the mechanism proposed in a different context (high levels of the H3K27me3-competing H3K37me3 mark) by Popovic et al. (2014): loss of accessibility to PRC2 binding to H3K27me3 would lead to an increase in nucleoplasmic free PRC2 concentration and greater recruitment to remaining binding sites. Our hypothesis is supported by the observation that many of the retention sites are found at CpG islands where PRC2 may be recruited by factors that may include histone-free linker DNA, MTF2, KDM2B-recruited PRC1 and JARID2 (Herz and Shilatifard, 2010; van Mierlo et al., 2019; Oksuz et al., 2018; Perino et al., 2019; Wang et al., 2017). The presence of H3K27me3 at these PRC2 accumulation sites could be due to incomplete inhibition of EZH2 activity, combined with the greater levels of PRC2. As our EZH2 inhibitors are 100 fold more selective toward EZH2 than to EZH1 (Bradley et al., 2014), it is also possible that the residual H3K27me3 in the cells after EZH2 inhibition at least in some part reflects the activities of EZH1-containing PRC2.

We observe a small peak of EZH2 or SUZ12 at TSSs in clusters 3 and 5, with actively transcribing genes. This is consistent with observations that PRC2 is recruited to nucleosome-free regions of chromatin (Wang et al., 2017), and models that suggest PRC2 “samples” open chromatin throughout the genome, only writing H3K27me3 at inactive genes. Our observation of loss of PRC2 at active genes suggests that recruitment or residence time at active TSSs is impaired by EZH2 inhibition.

Our observation that PRC2 accumulates preferentially at CpG islands upon EZH2 inhibition supports the hypothesis that these sites are the PRC2 nucleation sites in the genome. However, only 12.9% of the CpG islands are occupied by a retention site in EZH2 inhibitor treated KARPAS-422 cells, which may be explained by additional recruitment and retention requirements such as the state of DNA methylation and lack of gene transcription. Interestingly, our analysis showed that retention sites are better conserved among cell lines from the same tissue of origin, and are only moderately conserved between cell lines from different tissues. The observations point to a dynamic PRC2 nucleation site selection strategy; although PRC2 nucleates at CpG islands, different sets of CpG islands are utilized for PRC2 nucleation in different tissue types, reflecting the tissue-specific chromatin states and expression patterns that also influence recruitment.

### Up-regulation of bivalent genes is independent of sensitivity to EZH2 inhibitor

Given that H3K27me3 is known to maintain gene repression, we might expect that EZH2 inhibition would lead to up-regulation of H3K27me3-marked genes. While we do see some up-regulation of H3K27me3-marked genes, there are many up-regulated genes that do not have H3K27me3, and many H3K27me3-marked genes that are not up-regulated -- including some that are down-regulated. This broad range of up-regulation could be due to secondary effects of genes that are induced upon loss of H3K27me3, but could also be due to other mechanisms. We do see greater up-regulation of bivalent (H3K27me3 + H3K4me3) genes than genes having either mark alone, suggesting that some of the up-regulation is a direct effect of H3K27me3 reduction at poised genes. Down-regulation, on the other hand, is largely limited to genes that are not marked by H3K27me3, and is not explained by direct H3K27me3 targeting.

GCB-DLBCL often has activating mutations of EZH2, and/or high expression of EZH2. In the germinal center reaction required for normal immune function, there is rapid cell division accompanied by somatic hypermutation for antibody maturation; cells in the germinal center reaction are maintained in a proliferative state partly by PRC2-mediated repression of differentiation genes that lead to cellular differentiation into plasma and memory B cells (Béguelin et al., 2013 and references therein). Many of these genes have bivalent promoters carrying both H3K4me3 and H3K27me3, reflecting the transient nature of their silencing. It has been proposed that the PRC2 activating mutation in GCB-DLBCL hyper-methylates the promoters of these differentiation genes, making the activation of these genes unlikely and trapping the cells in an indefinite proliferative state. EZH2 inhibitor treatment leads to partial depression of many of the differentiation genes; however, as both sensitive and insensitive GCB-DLBCL cell lines demonstrate similar retention of H3K27me3 at differentiation genes, loss of H3K27me3 leading to direct up-regulation of expression does not fully explain the variation in sensitivity to EZH2 inhibitors.

## Acknowledgments

We thank Charlie Hatton for his help in data processing and computational infrastructure at Constellation. Constellation provided funding for materials and equipment to carry out the experiments and paid the publication fee. Work in the Tucker-Kellogg lab was supported by the Singapore Ministry of Education Academic Research Fund (AcRF) Tier 1 grant R154-000-693-112 to Greg Tucker-Kellogg.

## Author Contributions

The experimental work was conceived by PT and CCY. CCY designed and carried out the experimental work and conceived of the experimental/bioinformatics integration. Bioinformatic analyses and visualizations were designed and carried out by AJ, BB and GTK. Defining retention sites using chromatin state models was conceived by AJ. Work between Constellation and NUS was organised and coordinated by BB and GTK. CCY, AJ, GTK and BB wrote the initial drafts. All authors reviewed and edited the final manuscript.

## Methods

### Cell lines and antibodies

The H3K27me3 and H3K4me3 data was previously reported (Egan et al., 2016). KARPAS-422 cells were cultured and treated as previously described (Bradley et al., 2014). The following antibodies were used: H3K27me3 (Cell Signaling Technology 9730), H3K4me3 (Abcam ab8580), EZH2 (Millipore 07-689), SUZ12 (Active Motif 39357), total histone H3 (Cell Signaling Technology 3638), beta-Actin (Life Technologies AM4302).

### ChIP, ChIP-seq, and ChIP-seq data processing

Chromatin IP and ChIP-seq were carried out as previously described (Bradley et al., 2014). Custom primers used in ChIP-qPCR are: BHLHE41: F-acgtgcggaacgtaccat, R-cttctctcgccgccttct, UPL probe #52. KLHL29: R-ggtccttcagaggtgtggac, R-cgccctctttatgacactcc, UPL probe #15. RUNX2: F-tccaagctgcaaactcaagtc, R-ccggagtctttggaacacc, UPL probe #87. ABAT: F-gctgaactctttctctgccttta, R-gcgatcaccaagtcctcataa, UPL probe #58. MPEG1: F-gcctaacaggtagctatcctaattttt, R-gtgtggggtcgggatttt, UPL probe #20.

#### Normalization

For quantitative comparison of genome-wide mark patterns between DMSO and EZH2 inhibitor treated samples, *Drosophila* chromatin and *Drosophila* H2AV antibody (Active Motif 39715) were added to regular ChIP assays as described previously (Egan et al., 2016). For visualization, the total ChIP-seq signal was scaled to equalize the Drosophila component across samples for a given antibody (Egan et al., 2016). For data generated with fly chromatin spike-in, we created a combined human (hg19) and Drosophila (dm6) genome file from UCSC by combining the FASTA files [http://hgdownload.cse.ucsc.edu/goldenPath/hg19/bigZips/hg19.2bit and http://hgdownload.cse.ucsc.edu/goldenPath/dm6/bigZips/dm6.2bit]. Human chromosomes and Drosophila chromosomes were labeled differently to allow later separation. For ChIP-seq data generated without fly chromatin spike-in (H3K27me3 and EZH2 data for SU-DHL-6, KMS28BM, and KMS28PE, and EZH2 data for RPMI8226), we used normalization factors estimated based on western blot experiments. Only the human (hg19) genome file from UCSC was used for this data. Normalization factors for all ChIP-seq samples are listed in Table S2.

#### Mapping and coverage

FASTQ files containing 50-base-pair reads were mapped to either the combined human-fly genome or human genome using Bowtie 2 version 2.2.5 and compressed to BAM files using Samtools version 1.2; samtools was also used to sort and index the aligned read files. Duplicate reads were removed using the MarkDuplicates function in picard-tools, with lenient validation stringency. For reads mapped to the combined genome, mapped reads were then separated into reads mapped to human and fly chromosomes using the differential annotation used for the chromosomes. BamToBed from the Bedtools suite (version 2.24) converted the BAM files to intervals; bedtools’ slop function along with chromosome sizes obtained from UCSC was used to extend the reads to the 200bp fragment size.

#### Visualizations

Normalized BEDGRAPH and BIGWIG files were generated using the normalization factors; details are in the supplemental methods. For viewing in IGV (Robinson et al., 2011; Thorvaldsdóttir et al., 2013), BEDGRAPH files were converted to BIGWIG files using bedGraphToBigWig utility from UCSC. Within each IGV plot, y-axis scales are consistent across the genes for each mark. A heatmap for global mark levels was generated using ngsplot (Shen et al., 2014). The program was modified to use custom normalization factors. Clustering was done using K means clustering. Average profiles were obtained by ngsplot, again modified to use custom normalization factors, based on the coverage from BAM files. Scatter plots of change of ChIP-seq signal at TSS used log fold change as described in the supplemental methods.

### Chromatin state using ChromHMM

#### Model generation

Because the PRC2 components and H3K27me3 marks have broad and noisy signals, we chose to use chromatin states defined across multiple ChIP-seq tracks to assess chromatin state. We applied ChromHMM (version 1.0) (Ernst and Kellis, 2012) to the EZH2 and H3K27me3 tracks treated with DMSO or EZH2 inhibitor. Our code for running ChromHMM on our data is available upon request. H3K27me3 and EZH2 signal near TSSs showed patterns we were most interested in. Therefore we developed the ChromHMM models using ChIP-seq data within a region around TSSs starting 500 nucleotides upstream and going 2100 nucleotides into each gene. TSSs were defined using the transcription start site information from the hg19 refGene table from UCSC, created as described in supplemental methods. The txStart position was used for + strand genes, and the txEnd position was used for the - strand genes.

ChromHMM input files were generated from the BED files only containing isolated TSSs. From the original set of genes, we created BED files representing the range −500 to +2100 around each TSS. We filtered to keep only TSSs with non-overlapping regions; this step removed potentially confusing regions containing multiple TSSs. Minus stranded genes were flipped in order to minimize strand-bias. Each gene was taken as a separate chromosome so that segmentation by ChromHMM will occur at the same relative distance away from TSSs. These input files were binarized with BinarizeBed function then were used to generate a model using LearnModel function (with options -p 6 -s 123 -r 1000 -printposterior).

#### Model application

To apply the ChromHMM model genome-wide, we ran BinarizeBed on the full-genome ChIP-seq data. We then used MakeSegmentation (with option -printposterior) to apply the learned model to the genome-wide binarized data. The output segment file was filtered to keep only segments having posterior probability at least 0.8.

#### Retention site identification

Retention state was defined to be the state with high emission probability (at least 0.75) of either EZH2 or H3K27me3 mark upon EZH2 inhibition. All segments in the retention state were taken to be retention sites (loci). Retention genes were identified by finding genes with its −500 +2100 regions around TSS with retention sites.

#### Bivalent retention identification

To identify bivalent retention sites, the same pipeline was used with 8 input tracks (4 marks -- H3K27me3, EZH2, SUZ12, and H3K4me3 -- with and without EZH2 inhibition). Bivalent retention state(s) were defined as the state with high emission probability (at least 0.9) of both EZH2 and H3K4me3. All segments in the bivalent retention states were taken to be bivalent retention loci. Bivalent gene candidates were first identified by finding genes with its −500 +2100 regions around TSS with bivalent retention sites. This list of genes were compared to the retention genes list obtained earlier in the 2 marks analysis. The final list of bivalent genes were determined based on whether the bivalent gene candidate was also picked up by the earlier analysis as a retention gene. Emission probabilities of the retention states can be viewed in Table S1.

#### Finding overlap to enhancer sites, promoters, CpG islands

Retention sites overlapping various genomic regions were identified by intersecting the respective files using bedtools intersect (Quinlan, 2014). Fold enrichment was calculated based on the relative fraction of coverage by retention sites, using ChromHMM’s OverlapEnrichment function.

### RNA-seq data processing

KARPAS-422 cells treated with DMSO or EZH2 inhibitor were harvested and total RNA was isolated with Qiagen RNeasy Plus Mini kit (Qiagen 74136). Downstream sample processing, library generation and deep sequencing were carried out using the services of Ocean Ridge Biosciences (https://www.oceanridgebio.com/). RNA-seq data was processed in R/Bioconductor using library DESeq2. Gene features were obtained from TxDb.Hsapiens.UCSC.hg19.knownGene using exonsBy() with parameter by=”tx”. DESeq2 function summarizeOverlaps was run on the aligned BAM files using those exon features, Union mode, singleEnd=FALSE, ignore.strand=TRUE, and fragments=TRUE. DESeq was run on the dataset after specifying which samples were treated with EZH2i. The results were written to a CSV file for further analysis (Karpas_DESeq_output.R). Transcripts with base mean expression lower or equal to 9 were considered no expression. Transcripts with base mean expression higher than 9, but adjusted p-value higher than or equal to 0.1 were considered not differentially expressed. Transcripts with base mean expression higher than 9 and adjusted p-value lower than 0.1 were considered differentially expressed. Among them, if log2foldchange value was greater than 0, the transcripts were considered as upregulated. The rest were considered as downregulated.

### Signature analysis

Web-based signature analysis was performed the MSigDB “Investigate Gene Sets” tool (Subramanian et al., 2005) at the Broad Institute, with default parameters and the C2 (curated gene sets) category (Liberzon et al., 2011). In order to include additional gene sets, we downloaded the full gene sets from the Broad Institute, as well as supplemental data from the respective papers, and used the GeneOverlap (v.1.14.0) Bioconductor package (Shen, 2019) to calculate *p* values based on the Fisher’s exact test, as well as an odds ratio for strength of association. When plotted as heatmaps, *p* values were converted to false discovery rate estimates based on the total number of tests, and plotted for tests with FDR < 0.1 and odds ratio >= 2.

## Supplemental Information titles and legends

**Figure S1:** EZH2 inhibition results in genome-wide reduction of H3K27me3. (A) Western blots showing the global levels of EZH2, SUZ12, H3K27me3, and H3K4me3 in KARPAS-422 cells treated with DMSO or 1.5μM of CPI-169 for 4 and 8 days. While the total levels of EZH2, SUZ12, and H3K4me3 remain relatively unchanged, the level of H3K27me3 is reduced upon EZH2 inhibition. Anti-ACTB and anti-H3 immunoblots show constant levels of beta-Actin and Histone H3. (B) Normalized KARPAS-422 ChIP-seq tracks at three individual loci reflecting the global pattern of change after 8 days of CPI-360 treatment at 1.5μM CPI-360: levels of H3K27me3 decrease, while EZH2 and SUZ12 occupancy levels are relatively unchanged. (C) Average normalized KARPAS-422 ChIP-seq signal intensity for H3K27me3, H3K4me3, EZH2, and SUZ12 over all TSSs (+/− 1.5kb), either DMSO or CPI-360 treated. H3K27me3 level is reduced while EZH2, EED, and H3K4me3 levels show little change.

**Figure S2**: Scatter plot of 10,000 randomly sampled 1kb regions throughout the genome, showing strong correlation between H3K4me3 signal regardless of treatment.

**Figure S3**: Change in H3K27me3 vs change in EZH2 for each H3K27me3 cluster

**Figure S4**: Change in EZH2 vs baseline H3K27me3, with cluster 1 colored red showing high EZH2 retention

**Figure S5**: ChIP-qPCR of H3K27me3 and EZH2 levels before and after CPI-169 treatment. Three loci were selected from ChIP-seq data to be sites of retention of H3K27me3; the other two sites show loss of H3K27me3 by ChIP-seq. Error bars represent standard deviation from two independent experiments.

**Figure S6**. Expression changes for each H3K27me3 cluster from Figure 2A. (A) Top: Box and whiskers plot of natural log mean expression in each cluster for the different treatments. Middle: Violin plots of log2 fold change of gene expression. Clusters 1, 2 and 4 show more up-regulated genes than down-regulated genes, and clusters 3 and 5 show more down-regulated genes than up-regulated genes. Bottom: Pie charts showing fraction of genes that are up- or down-regulated in each cluster: no expression (white), no differential expression (gray), up-regulated (blue), down-regulated (red). The pie charts show that gene expression changes are not consistent within each cluster. (B) The table shows the number of genes in each pie chart. For numbers, see Table S3.

**Figure S7**. Retention sites across 6 cell lines used in this study. (A) Examples of retention sites showing that EZH2 accumulation is only partially conserved. MDGA1 is a retention gene in all cell lines, KLHL29 is retained in 5 out of 6 cell lines, while NR4A1 is only retained in KARPAS-422. (B) Genes that are retention genes in all 6 cell lines.

**Figure S8**. Analysis of set intersections from Venn diagrams of TSS retention site overlaps. (A) <u>Upset plot of TSS retention loci from all six cell lines</u>. All set intersections with set size greater than 15 loci are shown using the UpSet intersection method (Lex et al. 2014. “UpSet: Visualization of Intersecting Sets.” *IEEE Transactions on Visualization and Computer Graphics* 20 (12). IEEE: 1983–92). The size of the intersection is shown as a vertical bar, and the identity of the intersecting sets is shown by the connected dots below the bars. Horizontal bars (left) show the total number of retention loci in each cell line. (B) <u>Enrichment of H3K27me3 sites at retention loc</u>i were calculated for each intersection with a Fisher’s exact test, followed by a Benjamini-Hochberg correction based on the number of intersections tested. Each bar is shown as the estimate of the Fisher’s exact test (a form of enrichment over chance). Bars are shaded based on FDR < 0.05 (corrected for the number of distinct intersections tested). (C) <u>Enrichment of intersections over chance.</u> Chance intersections were simulated by taking random draws of the observed number of retention loci for six cell lines from a gene pool of the same size as the data. 100 such random gene sets were drawn, and intersection sizes calculated. The enrichment was calculated as (1 + observed intersection size)/(1 + max(chance intersection size)) to avoid dividing by zero.

**Table S1**. Emission probabilities of ChromHMM model used in retention site definition

**Table S2**. Normalization factors used for ChIP-seq

**Table S3**. Number of differentially expressed genes by cluster

**Table S4**. Number of differentially expressed genes by baseline histone mark

## Supplemental Methods

- Generation of normalized ChIP-seq coverage signal
- Transcription start site reference
- Scatter plots of ChIP-seq signal

## References

Béguelin, W., Popovic, R., Teater, M., Jiang, Y., Bunting, K.L.L., Rosen, M., Shen, H., Yang, S.N.N., Wang, L., Ezponda, T., et al. (2013). EZH2 Is Required for Germinal Center Formation and Somatic EZH2 Mutations Promote Lymphoid Transformation. Cancer Cell 23, 677–692.

Béguelin, W., Teater, M., Gearhart, M.D., Calvo Fern??ndez, M.T., Goldstein, R.L., Calvo Fernández, M.G., Hatzi, K., Rosen, M., Shen, H., Corcoran, C.M., et al. (2016). EZH2 and BCL6 Cooperate to Assemble CBX8-BCOR Complex to Repress Bivalent Promoters, Mediate Germinal Center Formation and Lymphomagenesis. Cancer Cell 30, 197–213.

Bernstein, B.E., Mikkelsen, T.S., Xie, X., Kamal, M., Huebert, D.J., Cuff, J., Fry, B., Meissner, A., Wernig, M., Plath, K., et al. (2006). A bivalent chromatin structure marks key developmental genes in embryonic stem cells. Cell 125, 315–326.

Bradley, W.D., Arora, S., Busby, J., Balasubramanian, S., Gehling, V.S., Nasveschuk, C.G., Vaswani, R.G., Yuan, C.C., Hatton, C., Zhao, F., et al. (2014). EZH2 inhibitor efficacy in non-Hodgkin’s lymphoma does not require suppression of H3K27 monomethylation. Chem. Biol. 21, 1463–1475.

Chan, K.M., Han, J., Fang, D., Gan, H., and Zhang, Z. (2013). A lesson learned from the H3.3K27M mutation found in pediatric glioma: a new approach to the study of the function of histone modifications in vivo? Cell Cycle Georget. Tex 12, 2546–2552.

Cooper, S., Dienstbier, M., Hassan, R., Schermelleh, L., Sharif, J., Blackledge, N.P., De Marco, V., Elderkin, S., Koseki, H., Klose, R., et al. (2014). Targeting polycomb to pericentric heterochromatin in embryonic stem cells reveals a role for H2AK119u1 in PRC2 recruitment. Cell Rep. 7, 1456–1470.

Di Croce, L., and Helin, K. (2013). Transcriptional regulation by Polycomb group proteins. Nat. Struct. Mol. Biol. 20, 1147–1155.

Egan, B., Yuan, C.-C., Craske, M.L., Labhart, P., Guler, G.D., Arnott, D., Maile, T.M., Busby, J., Henry, C., Kelly, T.K., et al. (2016). An Alternative Approach to ChIP-Seq Normalization Enables Detection of Genome-Wide Changes in Histone H3 Lysine 27 Trimethylation upon EZH2 Inhibition. PloS One 11, e0166438.

Ernst, J., and Kellis, M. (2012). ChromHMM: automating chromatin-state discovery and characterization. Nat. Methods 9, 215–216.

Garapaty-Rao, S., Nasveschuk, C., Gagnon, A., Chan, E.Y., Sandy, P., Busby, J., Balasubramanian, S., Campbell, R., Zhao, F., Bergeron, L., et al. (2013). Identification of EZH2 and EZH1 small molecule inhibitors with selective impact on diffuse large B cell lymphoma cell growth. Chem. Biol. 20, 1329–1339.

Girard, N., Bazille, C., Lhuissier, E., Benateau, H., Llombart-Bosch, A., Boumediene, K., and Bauge, C. (2014). 3-Deazaneplanocin A (DZNep), an inhibitor of the histone methyltransferase EZH2, induces apoptosis and reduces cell migration in chondrosarcoma cells. PloS One 9, e98176.

Gulati, N., Béguelin, W., and Giulino-Roth, L. (2018). Enhancer of zeste homolog 2 (EZH2) inhibitors. Leuk. Lymphoma 59, 1574–1585.

van Heeringen, S.J., Akkers, R.C., van Kruijsbergen, I., Arif, M.A., Hanssen, L.L.P., Sharifi, N., and Veenstra, G.J.C. (2014). Principles of nucleation of H3K27 methylation during embryonic development. Genome Res 24, 401–410.

Herz, H.-M., and Shilatifard, A. (2010). The JARID2–PRC2 duality. Genes Dev. 24, 857–861.

Hock, H. (2012). A complex Polycomb issue: The two faces of EZH2 in cancer. Genes Dev. 26, 751–755.

Italiano, A., Soria, J.-C., Toulmonde, M., Michot, J.-M., Lucchesi, C., Varga, A., Coindre, J.-M., Blakemore, S.J., Clawson, A., and Suttle, B. (2018). Tazemetostat, an EZH2 inhibitor, in relapsed or refractory B-cell non-Hodgkin lymphoma and advanced solid tumours: a first-in-human, open-label, phase 1 study. Lancet Oncol. 19, 649–659.

Kirmizis, A., Bartley, S.M., Kuzmichev, A., Margueron, R., Reinberg, D., Green, R., and Farnham, P.J. (2004). Silencing of human polycomb target genes is associated with methylation of histone H3 Lys 27. Genes Dev. 18, 1592–1605.

Ku, M., Koche, R.P., Rheinbay, E., Mendenhall, E.M., Endoh, M., Mikkelsen, T.S., Presser, A., Nusbaum, C., Xie, X., Chi, A.S., et al. (2008). Genomewide analysis of PRC1 and PRC2 occupancy identifies two classes of bivalent domains. PLoS Genet. 4.

Laugesen, A., Højfeldt, J.W., and Helin, K. (2019). Molecular Mechanisms Directing PRC2 Recruitment and H3K27 Methylation. Mol. Cell 74, 8–18.

Lenz, G., Wright, G.W., Emre, N.C.T., Kohlhammer, H., Dave, S.S., Davis, R.E., Carty, S., Lam, L.T., Shaffer, A.L., Xiao, W., et al. (2008). Molecular subtypes of diffuse large B-cell lymphoma arise by distinct genetic pathways. Proc. Natl. Acad. Sci. 105, 13520–13525.

Liberzon, A., Subramanian, A., Pinchback, R., Thorvaldsdóttir, H., Tamayo, P., and Mesirov, J.P. (2011). Molecular signatures database (MSigDB) 3.0. Bioinformatics 27, 1739–1740.

Malouf, G.G., Taube, J.H., Lu, Y., Roysarkar, T., Panjarian, S., Estecio, M.R., Jelinek, J., Yamazaki, J., Raynal, N.J., and Long, H. (2013). Architecture of epigenetic reprogramming following Twist1-mediated epithelial-mesenchymal transition. Genome Biol. 14, R144.

Margueron, R., and Reinberg, D. (2011). The Polycomb complex PRC2 and its mark in life. Nature 469, 343–349.

McCabe, M.T., Ott, H.M., Ganji, G., Korenchuk, S., Thompson, C., Van Aller, G.S., Liu, Y., Graves, A.P., Diaz, E., and LaFrance, L.V. (2012). EZH2 inhibition as a therapeutic strategy for lymphoma with EZH2-activating mutations. Nature 492, 108.

van Mierlo, G., Veenstra, G.J.C., Vermeulen, M., and Marks, H. (2019). The Complexity of PRC2 Subcomplexes. Trends Cell Biol.

Morin, R.D., Johnson, N.A., Severson, T.M., Mungall, A.J., An, J., Goya, R., Paul, J.E., Boyle, M., Woolcock, B.W., Kuchenbauer, F., et al. (2010). Somatic mutations altering EZH2 (Tyr641) in follicular and diffuse large B-cell lymphomas of germinal-center origin. Nat. Genet. 42, 181–185.

Oksuz, O., Narendra, V., Lee, C.-H., Descostes, N., LeRoy, G., Raviram, R., Blumenberg, L., Karch, K., Rocha, P.P., and Garcia, B.A. (2018). Capturing the onset of PRC2-mediated repressive domain formation. Mol. Cell 70, 1149–1162.

Perino, M., van Mierlo, G., Wardle, S.M., Marks, H., and Veenstra, G.J.C. (2019). Two distinct functional axes of positive feedback-enforced PRC2 recruitment in mouse embryonic stem cells. BioRxiv 669960.

Popovic, R., Martinez-Garcia, E., Giannopoulou, E.G., Zhang, Q., Zhang, Q., Ezponda, T., Shah, M.Y., Zheng, Y., Will, C.M., and Small, E.C. (2014). Histone methyltransferase MMSET/NSD2 alters EZH2 binding and reprograms the myeloma epigenome through global and focal changes in H3K36 and H3K27 methylation. PLoS Genet. 10, e1004566.

Quinlan, A.R. (2014). BEDTools: The Swiss-Army Tool for Genome Feature Analysis. Curr. Protoc. Bioinforma. 47, 11.12.1–34.

Robinson, J.T., Thorvaldsdóttir, H., Winckler, W., Guttman, M., Lander, E.S., Getz, G., and Mesirov, J.P. (2011). Integrative genomics viewer. Nat. Biotechnol. 29, 24.

Shen, L. (2019). GeneOverlap: Test and visualize gene overlaps.

Shen, L., Shao, N., Liu, X., and Nestler, E. (2014). ngs. plot: Quick mining and visualization of next-generation sequencing data by integrating genomic databases. BMC Genomics 15, 284.

Subramanian, A., Tamayo, P., Mootha, V.K., Mukherjee, S., Ebert, B.L., Gillette, M.A., Paulovich, A., Pomeroy, S.L., Golub, T.R., Lander, E.S., et al. (2005). Gene set enrichment analysis: a knowledge-based approach for interpreting genome-wide expression profiles. Proc Natl Acad Sci U A 102, 15545–15550.

Thorvaldsdóttir, H., Robinson, J.T., and Mesirov, J.P. (2013). Integrative Genomics Viewer (IGV): high-performance genomics data visualization and exploration. Brief Bioinform 14, 178–192.

Tiffen, J.C., Gunatilake, D., Gallagher, S.J., Gowrishankar, K., Heinemann, A., Cullinane, C., Dutton-Regester, K., Pupo, G.M., Strbenac, D., Yang, J.Y., et al. (2015). Targeting activating mutations of EZH2 leads to potent cell growth inhibition in human melanoma by derepression of tumor suppressor genes. Oncotarget 6, 27023–27036.

Wang, X., Paucek, R.D., Gooding, A.R., Brown, Z.Z., Ge, E.J., Muir, T.W., and Cech, T.R. (2017). Molecular analysis of PRC2 recruitment to DNA in chromatin and its inhibition by RNA. Nat Struct Mol Biol 24, 1028–1038.

Xu, B., On, D.M., Ma, A., Parton, T., Konze, K.D., Pattenden, S.G., Allison, D.F., Cai, L., Rockowitz, S., Liu, S., et al. (2015). Selective inhibition of EZH2 and EZH1 enzymatic activity by a small molecule suppresses MLL-rearranged leukemia. Blood 125, 346–357.

Yap, T.A., Winter, J.N., Leonard, J.P., Ribrag, V., Constantinidou, A., Giulino-Roth, L., Michot, J.-M., Khan, T.A., Horner, T., Carver, J., et al. (2016). A Phase I Study of GSK2816126, an Enhancer of Zeste Homolog 2(EZH2) Inhibitor, in Patients (pts) with Relapsed/Refractory Diffuse Large B-Cell Lymphoma (DLBCL), Other Non-Hodgkin Lymphomas (NHL), Transformed Follicular Lymphoma (tFL), Solid Tumors and Multiple Myeloma (MM). Blood 128, 4203–4203.

Zeng, D., Liu, M., and Pan, J. (2017). Blocking EZH2 methylation transferase activity by GSK126 decreases stem cell-like myeloma cells. Oncotarget 8, 3396–3411.

