## Supplemental Information for "EZH2 inhibition results in genome-wide PRC2 redistribution"

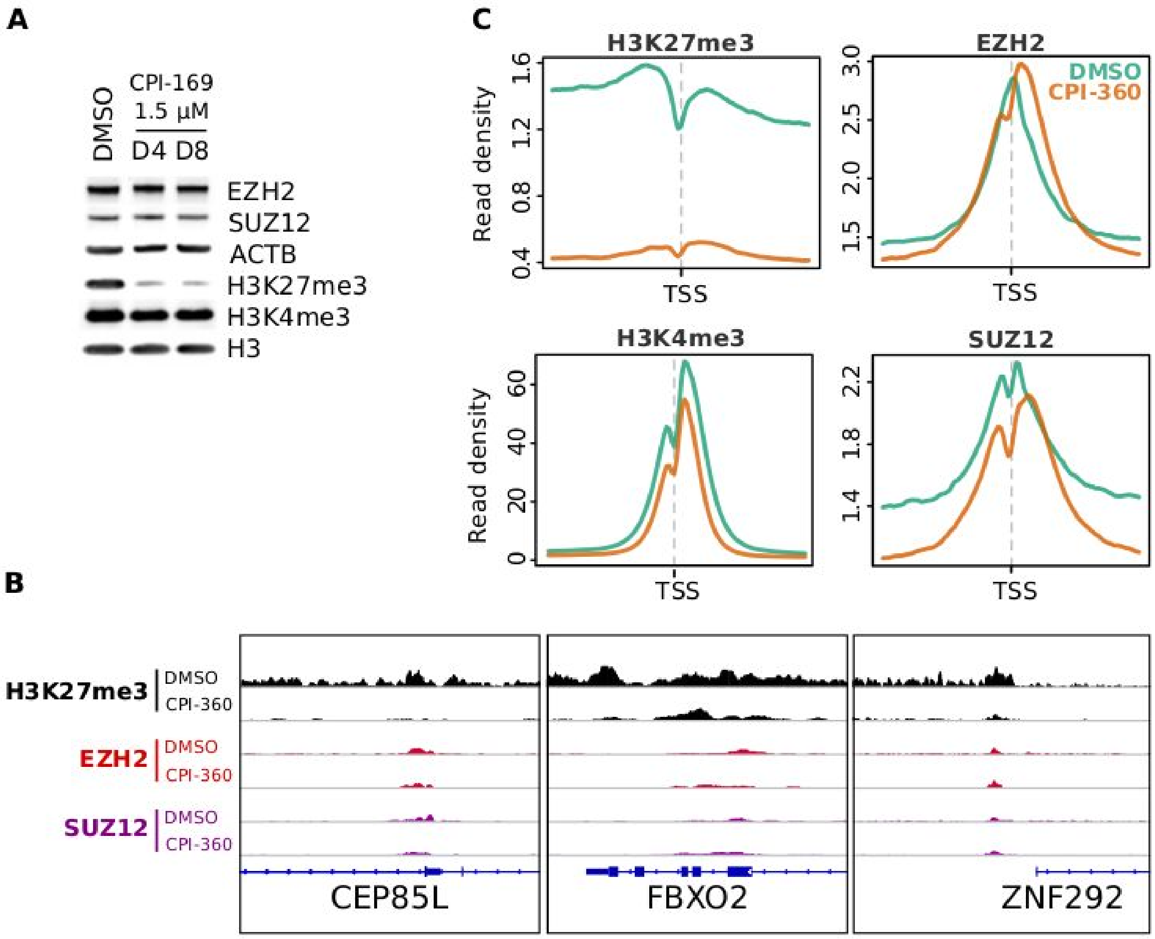


**Figure S1. EZH2 inhibition results in genome-wide reduction of H3K27me3**

A. Western blots showing the global levels of EZH2, SUZ12, H3K27me3, and H3K4me3 in KARPAS-422 cells treated with DMSO or 1.5µM of CPI-169 for 4 and 8 days. While the total levels of EZH2, SUZ12, and H3K4me3 remain relatively unchanged, the level of H3K27me3 is reduced upon EZH2 inhibition. Anti-ACTB and anti-H3 immunoblots show constant levels of beta-Actin and Histone H3.
B. Normalized KARPAS-422 ChIP-seq tracks at three individual loci reflecting the global pattern of change after 8 days of CPI-360 treatment at 1.5µM CPI-360: levels of H3K27me3 decrease, while EZH2 and SUZ12 occupancy levels are relatively unchanged.
C. Average normalized KARPAS-422 ChIP-seq signal intensity for H3K27me3, H3K4me3, EZH2, and SUZ12 over all TSSs (+/- 1.5kb), either DMSO or CPI-360 treated. H3K27me3 level is reduced while EZH2, EED, and H3K4me3 levels show little change.

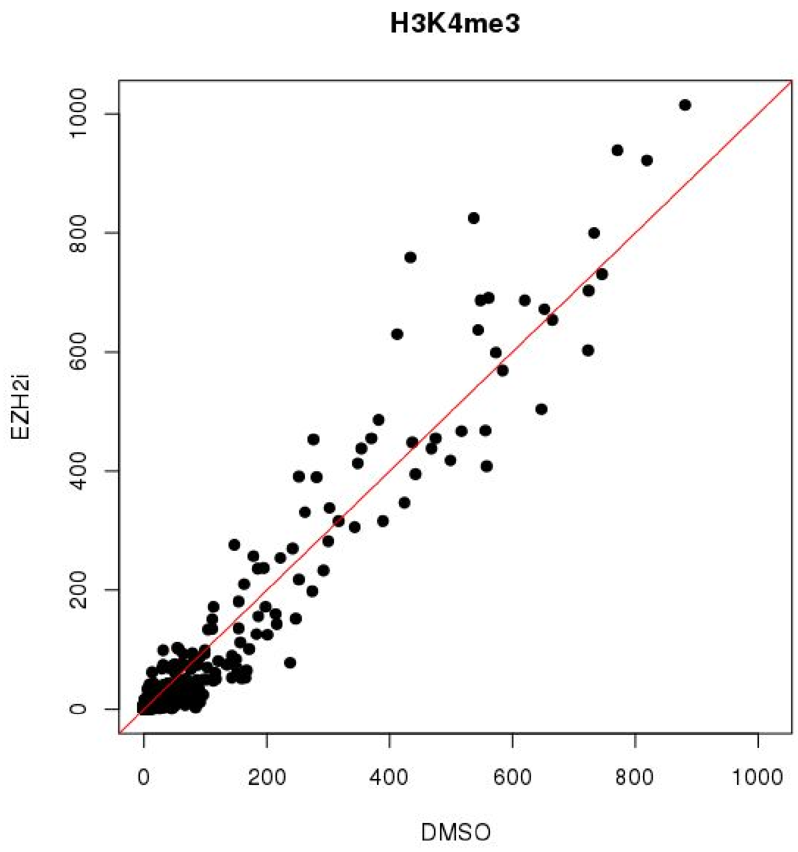


**Figure S2**

Scatter plot of 10,000 randomly sampled 1kb regions throughout the genome, showing strong correlation between H3K4me3 signal regardless of treatment.

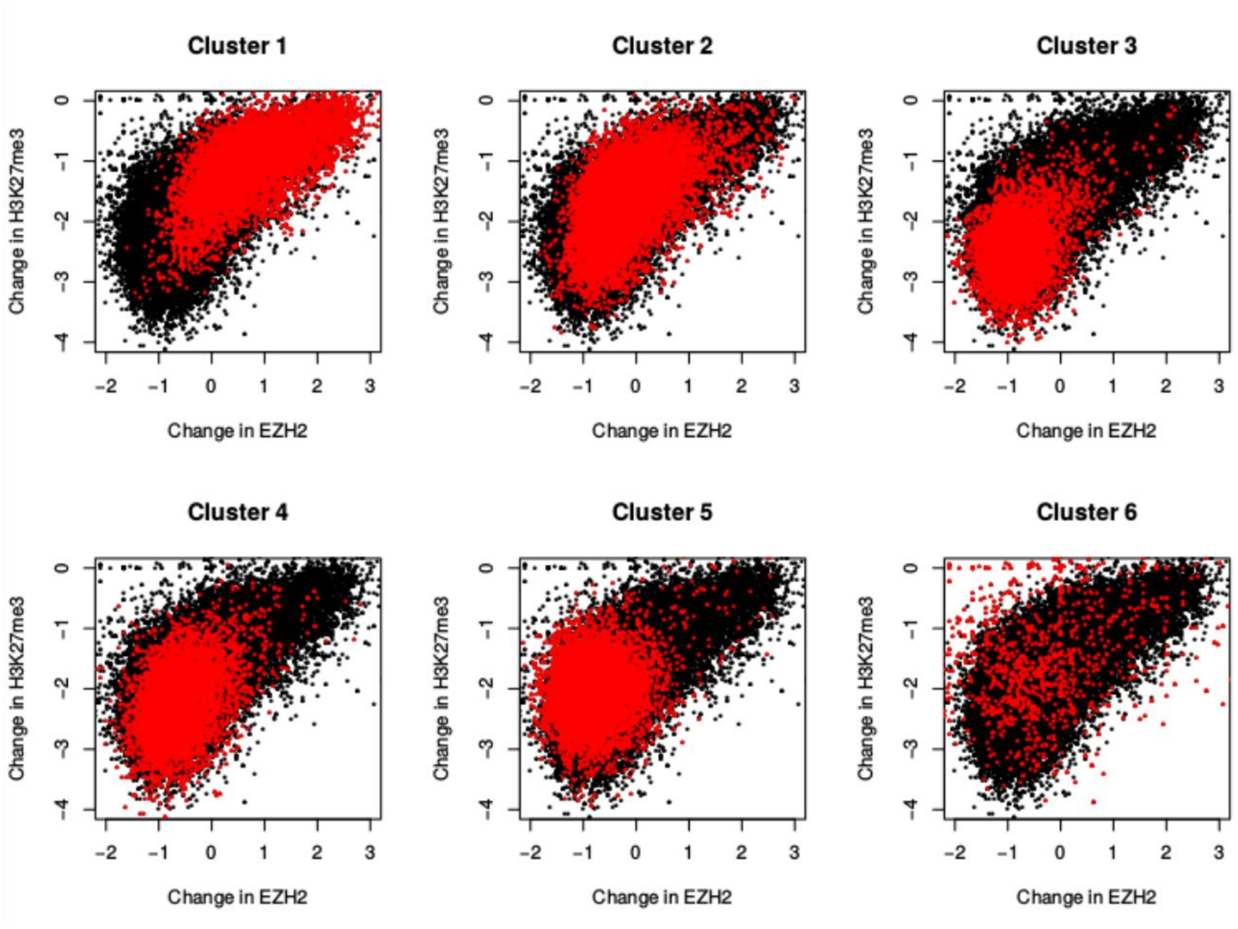


**Figure S3**

Change in H3K27me3 vs change in EZH2 for each H3K27me3 cluster

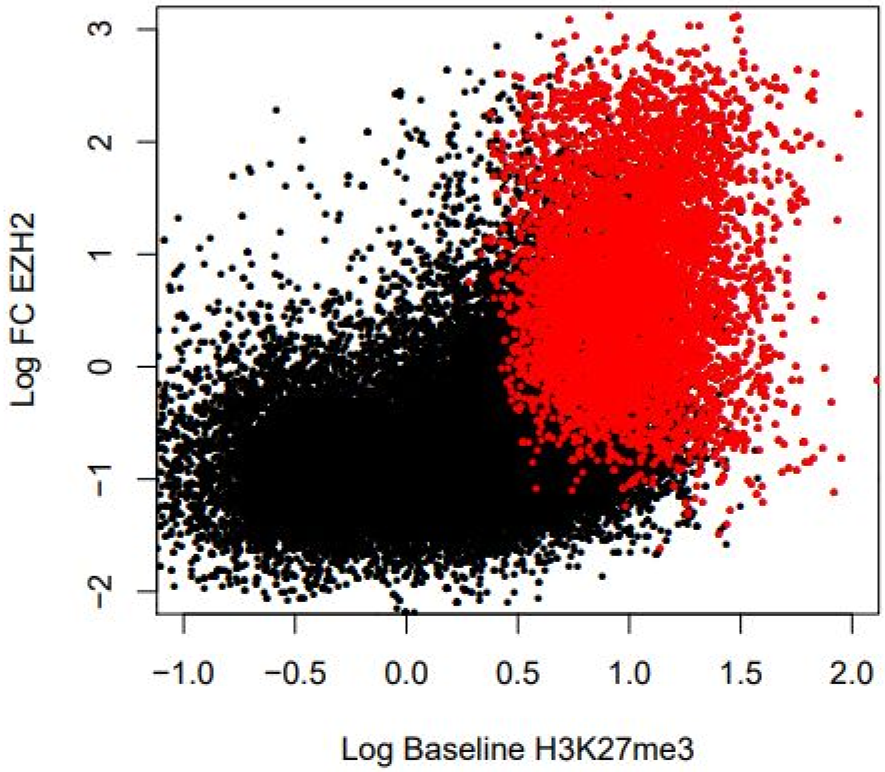


**Figure S4**

Change in EZH2 vs baseline H3K27me3, with cluster 1 colored red showing high EZH2 retention

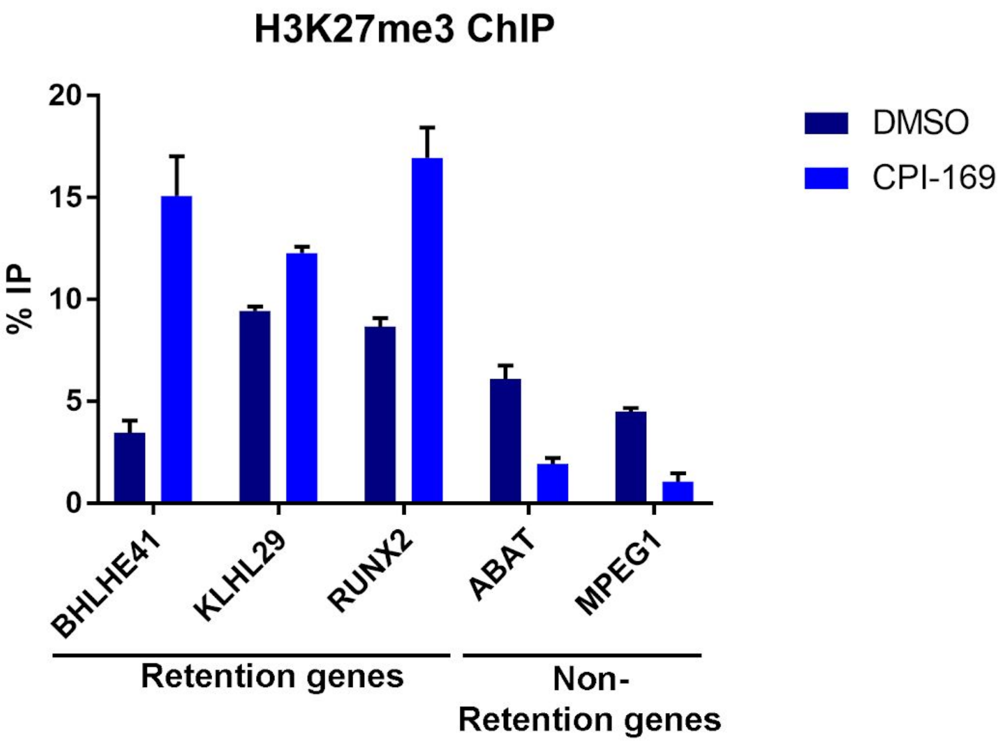


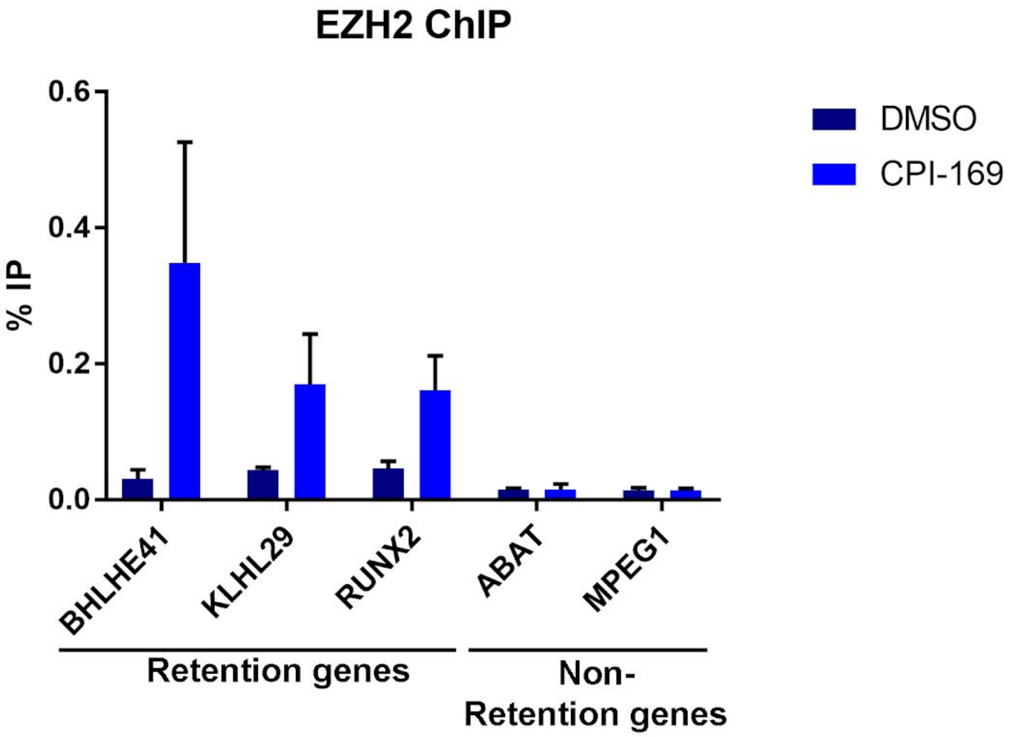


**Figure S5**

ChIP-qPCR of H3K27me3 and EZH2 levels before and after CPI-169 treatment. Three loci were selected from ChIP-seq data to be sites of retention of H3K27me3; the other two sites show loss of H3K27me3 by ChIP-seq. Error bars represent standard deviation from two independent experiments.

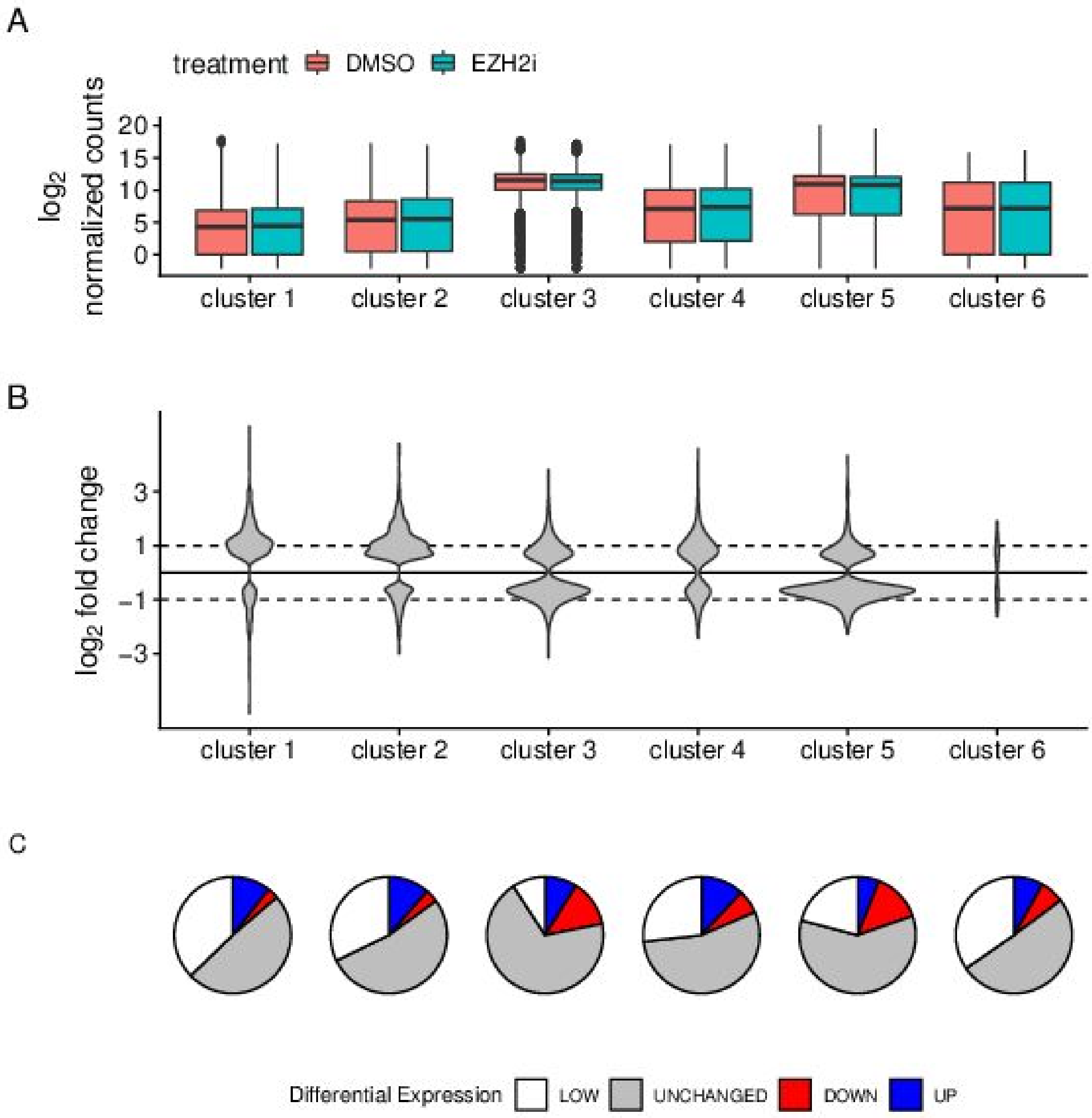


**Figure S6. Expression changes for each H3K27me3 cluster from Figure 2A.**

A. Top: Box and whiskers plot of natural log mean expression in each cluster for the different treatments. Middle: Violin plots of log2 fold change of gene expression. Clusters 1, 2 and 4 show more up-regulated genes than down-regulated genes, and clusters 3 and 5 show more down-regulated genes than up-regulated genes. Bottom: Pie charts showing fraction of genes that are up- or down-regulated in each cluster: no expression (white), no differential expression (gray), up-regulated (blue), down-regulated (red). The pie charts show that gene expression changes are not consistent within each cluster.

B. The table shows the number of genes in each pie chart. For numbers, see Table S3.

**A**

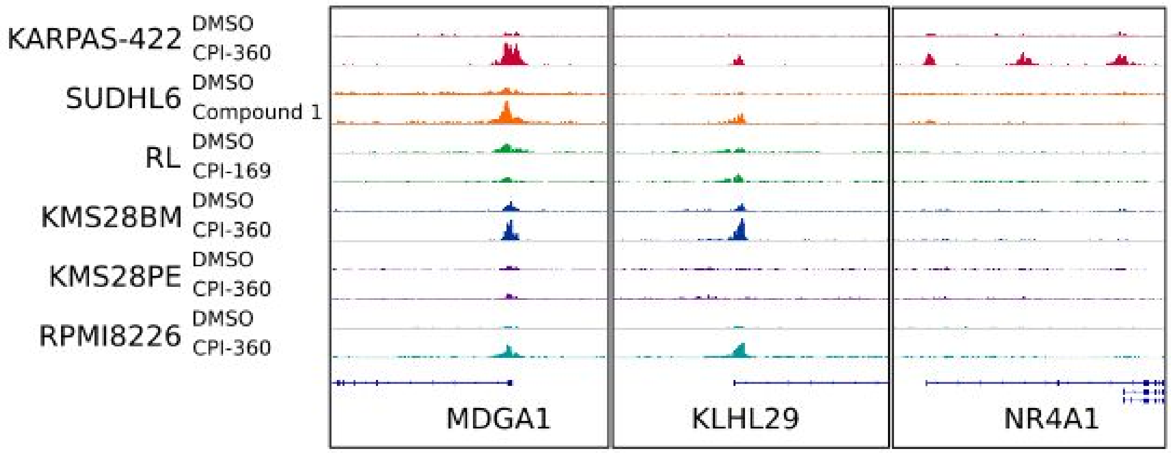


**B**

| | | | | |
|---|---|---|---|---|
| AMZ1 | GSX2 | MDGA1 | REXO1L2P | SMAD6 |
| C2CD4D | HES3 | MFSD4 | RHBDL3 | SMARCD3 |
| CDH22 | HES5 | MIR9-1 | RSPO3 | SOX9 |
| CHST1 | HOXB9 | NDRG4 | RTN4RL1 | SOX9-AS1 |
| CRLF1 | HOXC8 | NEUROD2 | RUNX2 | ST6GALNAC2 |
| DLL1 | HOXC-AS2 | OROF | SAMD11 | TENM4 |
| DUSP15 | INTS4P2 | P3H3 | SAMD14 | TMEM151B |
| FAM120B | KCNN1 | PAX2 | SEMA5B | TMEM59L |
| FAM131B | LOC100132111 | PDE4DIP | SERTAD4 | TRABD2A |
| FGFR1 | LOR | PTH1R | SERTAD4-AS1 | TTLL9 |
| GRIN2C | LRTM2 | RASSF5 | SLC30A3 | ZNF423 |

**Figure S7. Retention sites across 6 cell lines used in this study**

A. Examples of retention sites showing that EZH2 accumulation is only partially conserved. MDGA1 is a retention gene in all cell lines, KLHL29 is retained in 5 out of 6 cell lines, while NR4A1 is only retained in KARPAS-422.
B. Genes that are retention genes in all 6 cell lines.

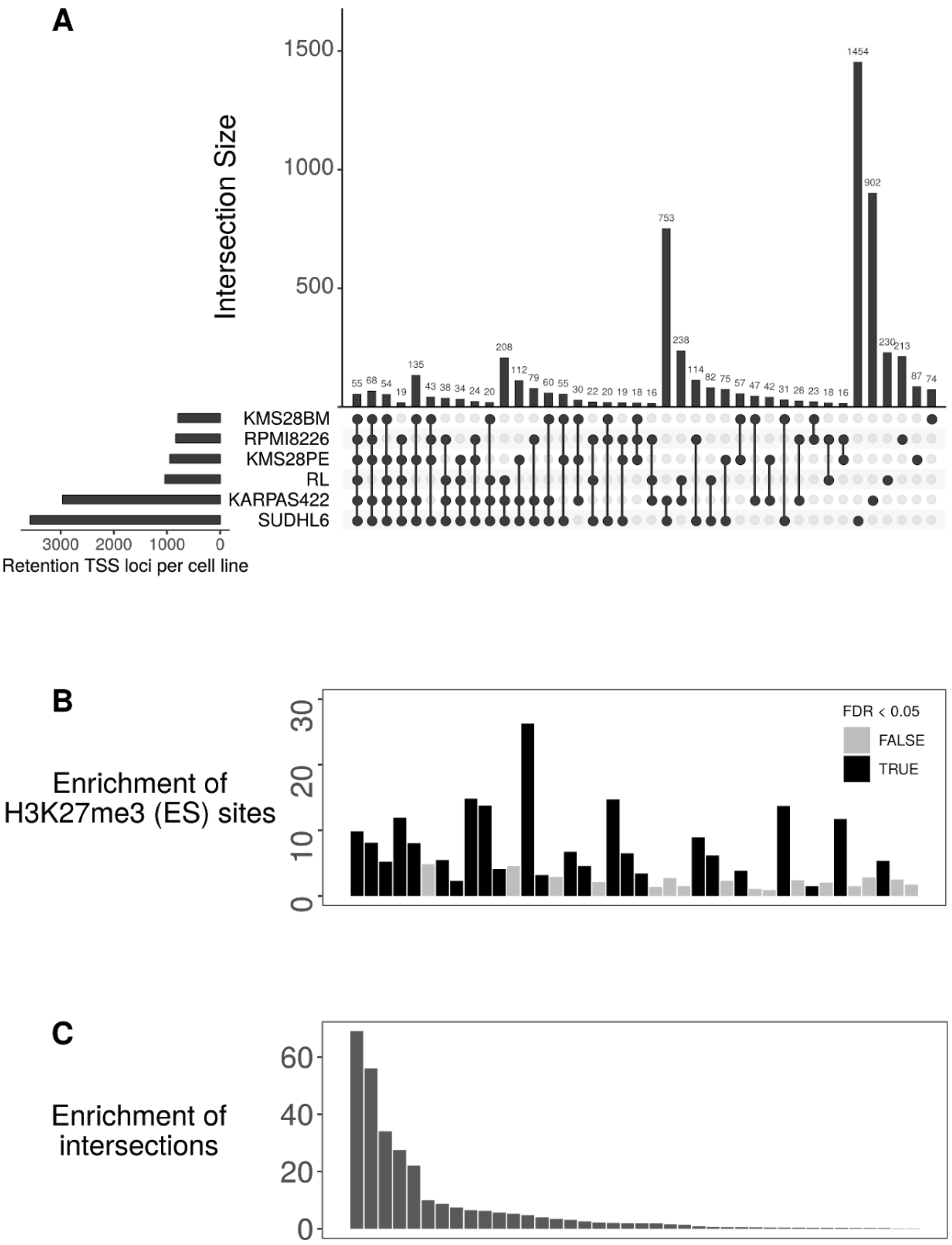


**Figure S8. Analysis of set intersections from Venn diagrams of TSS retention site overlaps.**

A. Upset plot of TSS retention loci from all six cell lines. All set intersections with set size greater than 15 loci are shown using the UpSet intersection method (Lex et al. 2014. “UpSet: Visualization of Intersecting Sets.” *IEEE Transactions on Visualization and Computer Graphics* 20 (12). IEEE: 1983–92). The size of the intersection is shown as a vertical bar, and the identity of the intersecting sets is shown by the connected dots below the bars. Horizontal bars (left) show the total number of retention loci in each cell line
B. Enrichment of H3K27me3 sites at retention loci were calculated for each intersection with a Fisher’s exact test, followed by a Benjamini-Hochberg correction based on the number of intersections tested. Each bar is shown as the estimate of the Fisher’s exact test (a form of enrichment over chance). Bars are shaded based on FDR < 0.05 (corrected for the number of distinct intersections tested).
C. Enrichment of intersections over chance. Chance intersections were simulated by taking random draws of the observed number of retention loci for six cell lines from a gene pool of the same size as the data. 100 such random gene sets were drawn, and intersection sizes calculated. The enrichment was calculated as (1 + observed intersection size)/(1 + max(chance intersection size)) to avoid dividing by zero.

**Table S1: Emission probabilities of ChromHMM model used in retention site definition**

| state (Emission order) | DMSO_EZH2 | DMSO_H3K27me3 | EZH2i_H3K27me3 | EZH2i_EZH2 |
|---|---|---|---|---|
| 1 | 0.1807081601 | 0.2112651928 | 0.09421693692 | 0.0215156805 |
| 2 | 0.002138122801 | 0.002630309496 | 0.001265338666 | 8.90E-12 |
| 3 | 8.81E-04 | 0.07978190847 | 0.1737998192 | 0.03987528863 |
| 4 | 0.1364802703 | 0.1906853959 | 0.5117773678 | 0.9025871541 |

**Table S2: Normalization factors used for ChIP-seq**

| **cell line** | **condition** | **antibody** | **normalization method** | **normalization factor** |
|---|---|---|---|---|
| SU-DHL-6 | DMSO | H3K27me3 | WBF | 1 |
| SU-DHL-6 | EZH2i | H3K27me3 | WBF | 0.3031984248 |
| KMS28BM | DMSO | H3K27me3 | WBF | 1 |
| KMS28BM | EZH2i | H3K27me3 | WBF | 0.1567738013 |
| KMS28PE | DMSO | H3K27me3 | WBF | 1 |
| KMS28PE | EZH2i | H3K27me3 | WBF | 0.5746972487 |
| RPMI8226 | DMSO | H3K27me3 | fly | 0.152146 |
| RL | DMSO | H3K27me3 | fly | 0.50771 |
| RL | EZH2i | H3K27me3 | fly | 0.04327 |
| RL | DMSO | EZH2 | fly | 0.24613 |
| RL | EZH2i | EZH2 | fly | 0.26586 |

**Table S3. Number of differentially expressed genes by cluster**

| | cluster 1 | cluster 2 | cluster 3 | cluster 4 | cluster 5 | cluster 6 |
|---|---|---|---|---|---|---|
| No expression | 1581 | 1733 | 396 | 872 | 1269 | 161 |
| Unchanged | 2082 | 2843 | 2975 | 1783 | 3604 | 235 |
| Upregulated | 454 | 617 | 383 | 391 | 362 | 38 |
| Downregulated | 132 | 194 | 562 | 213 | 831 | 31 |
| Total | 4249 | 5387 | 4316 | 3259 | 6066 | 465 |

**Table S4. Number of differentially expressed genes by baseline histone mark**

| | H3K27me3 | H3K27me3/H3K4me3 | H3K4me3 | unmarked | Total |
|---|---|---|---|---|---|
| Low/no expression | 1974 | 244 | 865 | 2929 | 6012 |
| Unchanged | 2355 | 1364 | 6492 | 3311 | 13522 |
| Downregulated | 114 | 210 | 1386 | 253 | 1963 |
| Upregulated | 353 | 776 | 836 | 280 | 2245 |
| Total | 4796 | 2594 | 9579 | 6773 | |

### Supplemental Methods

#### Generation of normalized ChIP-seq coverage signal

Using bedtools genomecov, BED format interval files were converted to coverage files in BEDGRAPH format (unnormalized coverage file). The average coverage of the unnormalized coverage file was calculated, assuming an effective genome length of 2,700,000,000 base pairs. Both DMSO and EZH2-inhibitor-treated samples were normalized to an average coverage signal of 1.0. EZH2-inhibitor-treated samples were then further normalized using a method that depended on whether fly spike-in was used. For samples without fly spike-in, the normalization factors were calculated as follows:

WB_ratio = (EZH2i signal from western blot) / (DMSO signal from western blot)
DMSO_dfactor = 1 / (average coverage of unnormalized DMSO coverage file)
EZH2i_dfactor = 1 / (average coverage of unnormalized EZH2i coverage file)
EZH2i_cfactor = EZH2i_dfactor * WB_ratio

For samples spiked with fly chromatin, the number of reads in the fly DMSO sample and fly EZH2-inhibitor-treated sample were determined using samtools flagstat. The normalization were calculated as follows:

DMSO_dfactor = 1 / (average coverage of unnormalized DMSO coverage file)
EZH2i_sfactor = Number of reads in the fly DMSO sample divided by the number of reads in the fly EZH2i-treated sample.
EZH2i_cfactor = DMSO_dfactor * EZH2i_sfactor

Normalized coverage files in BEDGRAPH were then generated using bedtools genomecov and with DMSO_dfactor and EZH2i_cfactor as scale factors for DMSO and EZH2-inhibitor-treated samples. For viewing in IGV (Thorvaldsdóttir et al. 2013, Robinson et al. 2011), BEDGRAPH files were converted to BIGWIG files using bedGraphToBigWig utility from UCSC.

The manipulations above equate to calculating the relative amount of global signal in treated vs control samples as (HT * FC/FT) / HC, where
HT = number of reads in the treated sample mapped uniquely to the human genome
HC = number of reads in the control sample mapped uniquely to the human genome
FT = number of reads in the treated sample mapped uniquely to the Drosophila genome
FC = number of reads in the control sample mapped uniquely to the Drosophila genome

### Transcription start site reference

Transcription start sites (TSSs) were defined using the hg19 refGene table from UCSC, created from the URL: http://genome.ucsc.edu/cgi-bin/hgTrackUi?db=hg19&g=refGene; (Pruitt et al. 2012). The table was obtained with the unix command “mysql --user=genome --host-genome-mysql.cse.ucsc.edu -A -D hg19 -e ‘select * from refGene’”.

### Scatter plots of ChIP-seq signal

Figures 2D and S2 plot change in H3K27me3 vs change in EZH2. Change in signal was calculated as log2((treated + 0.01)/(control + 0.01)). Figure 5B shows baseline H3K4me3 plotted against baseline H3K27me3. In both cases, we show the log based 2 of the (x+0.1), where x is the ChIP-seq signal.
